# Identifying spawning activity in aquatic species based on environmental DNA spikes

**DOI:** 10.1101/2020.01.28.924167

**Authors:** Satsuki Tsuji, Naoki Shibata

## Abstract

An understanding of the reproductive biology of aquatic organisms is crucial for the efficient conservation and management of species and/or populations. Nevertheless, conventional spawning surveys such as visual- and capture-based monitoring generally require laborious, time-consuming work and are subject to monitoring biases such as observer bias, as well as miscounts due to false spawning. In addition, direct capture often damages eggs or individuals. Thus, an efficient non-invasive approach for monitoring spawning activity on aquatic species would be a valuable tool to understand their reproductive biology and conserving biodiversity. Here, we proposed an environmental DNA (eDNA)-based approach for monitoring and understanding spawning activity by observing spikes in eDNA concentration after spawning activity. We found in cross experiments using two medaka species (*Oryzias latipes* and *O. sakaizumii*, 1:1 individual per tank) that an eDNA spike occurred in only male species after spawning activity. In addition, the magnitude of the eDNA spike was dependent on the number of spawning activities with egg and sperm release. In the field survey during the reproductive season, eDNA concentration after spawning were 3–25 times higher than before spawning. On the other hand, there was no increase in eDNA concentration during the non-reproductive season. Therefore, our results demonstrated that spikes in the eDNA concentration are mainly caused by the release of sperm during spawning activity, and it can be used as evidence of spawning in field survey. The presented approach could be a practical tool for studying reproductive biology and provides an opportunity to design effective conservation and environmental management actions.

## Introduction

Reproductive events are important parts of an aquatic organisms annual life history, and especially for rare species and important fishery target species, the understanding of spawning timing and location is critical for efficient conservation and management of species and/or populations (Danylchuk et al., 2011; Spear et al., 2015). Nevertheless, conventional spawning surveys (by direct observation of individuals or eggs) generally require laborious, time-consuming work and are subject to monitoring biases such as observer bias, as well as geographical constraints and miscounts due to false spawning (Caswell et al., 2004; Miller et al., 2012; Ko et al., 2013; Koster et al., 2013; Diana et al., 2015; Bylemans et al., 2016). In addition, particularly for rare and endangered species, traditional capture survey methods may threaten the persistence of species or populations because of additional mortality among the spawning population and/or eggs (Tsukamoto 2006; Wei et al., 2009; Engstedt et al., 2014). Thus, an efficient, noninvasive method of monitoring spawning activity would be a valuable tool to increase our understanding of species’ reproductive biology and conserving aquatic biodiversity.

Environmental DNA (eDNA) analysis has been used to estimate species distribution, and recent works have shown a relatively higher eDNA concentration in water during the reproductive season (Spear et al., 2015; Bracken et al., 2018; Tillotson et al., 2018; Takahashi et al., 2018; Thalinger et al., 2019). eDNA is a generic term used to refer to the DNA that has been shed by organisms into the surrounding environment, and it is thought to derive from skin, feces, mucus and reproductive materials (e.g. oocytes, ovarian fluid and sperm) (Ficetola et al., 2008; Merkes et al., 2014; Barnes and Turner. 2016; Takeuchi et al., 2019). Thus, in the case of externally-fertilizing species, an increase of eDNA concentration in the reproductive season would be expected. Furthermore, eDNA analysis could, potentially, avoid the restrictions associated with conventional spawning observations imposed by time, labour, monitoring biases and invasiveness because eDNA analysis requires only water sampling at the study site. Therefore, the monitoring for eDNA concentration changes would provide promising opportunities to efficiently and noninvasively monitor spawning activity in the area.

However, it is important to note that the seasonal increase of eDNA concentration is not always evidence of spawning; it can be caused by the formation of spawning aggregations (Takeuchi et al., 2019). While still controversial because eDNA persistence and release rates are not constant across different species and environmental conditions, many previous studies have shown a significant positive relationship between abundance/biomass of species and eDNA concentration (e.g., Takahara et al., 2012; Yamamoto et al., 2016; Doi et al., 2017). To obtain evidence of spawning activity, Bylemans et al. (2016) proposed an approach focused on a change in ratio of nuclear eDNA fragments to mitochondrial eDNA fragments during spawning. The increasing of the nuclear eDNA fragment after spawning probably caused by containing the released sperm which has more highly condensed and protected nuclear DNA than somatic cells as the eDNA source. Although this approach is sound, it is of limited use because it requires nuclear DNA sequence data, which are not well archived compared with those of mitochondrial DNA for invertebrates and considered to be lacking in variability at the genus or species level (Minamoto et al., 2017). Therefore, proposing a more versatile eDNA approach to monitor spawning activity would be extremely valuable.

Here, we propose a novel approach to monitoring and understanding spawning activity by observing the eDNA concentration spike after spawning activity. As suggested in previous studies (Bylemans et al., 2016; Takeuchi et al., 2019), when spawning occurs, the released sperm and/or oocytes and ovarian fluid are likely to become major sources of eDNA. These substances are mass released from individuals in a fraction of a second during spawning, and eDNA concentration should be shown a rapid increase. Thus, first, we examined which of factors (sperm release or oocytes and ovarian fluid release) causes an increase in eDNA concentration during spawning activities. In addition, we analysed the relationship between the magnitude of eDNA spike and the number of spawning activities with egg and sperm release. Finally, we examined whether we could detect the spawning activity of two medaka species in the field conditions based on the observation of the eDNA concentration spike.

## Materials and Methods

### 1. Study species and its spawning activity

The two Japanese medaka species, *Oryzias latipes* and *Oryzias sakaizumii*, are small (3–4 cm in standard length) freshwater fish (Iguchi et al., 2018) and listed as “Vulnerable” on the Red List of Threatened Species of Japan (Ministry of the Environment, Japan, 2015). However, many genetically distinct medaka varieties have been created and maintained by fish farmers because they can breed easily in captivity and are very popular as aquarium fish (Nakao et al., 2015). While both wild medaka species have black–grey body colouration and are morphologically similar, the most popular cultured medaka, originated from *O. latipes*, has orange–red body coloration. In this study, to identify the two medaka species easily, the orange–red variety of *O. latipes* and the wild type of *O. sakaizumii* were used for tank experiments. There is no difference in spawning activity between the orange–red medaka and wild type medaka, and they easily mate in both artificial and natural conditions (Sakaizumi et al., 1992; Kobayashi et al., 2012; Nakao et al., 2015).

Spawning activity of two medaka species occurs when the light condition, water temperature, and adequate nutritional status are at optimum. If they are kept under appropriate conditions, daily spawning can be induced any point in the year for up to three months’ duration. Optimum conditions for daily spawning are as follows: 14 hr of daylight and 10 hr of darkness, water temperature of 25–28°C, and feeding at least three times a day. Spawning activity in medaka species is usually observed within the first hour after the light is turned on (Kinoshita et al., 2009). The main steps are as follows: (1) following and positioning (the male approaches the female but keeps near distance); (2) dancing (the male swims in a circle in front of the female); (3) wrapping (the male wraps the female with his dorsal and anal fins); (4) egg release and sperm release; (5) separation (the male leaves the female); and (6) egg stripping (the attachment of the eggs to a substance). However, sometimes step 4, egg release and sperm release, does not occur, despite the warm-up steps (steps 1–3), and the female separates from the male (false spawning). It is difficult to distinguish true spawning from false spawning when observing from above.

Under field conditions, spawning activity in the medaka species can be observed from approximately May to September (daily mean water temperature > 20°C and day length > 12 hr). In a wild population, spawning activity starts from approximately 1 hr before sunrise and continues for approximately 6 hr (Kobayashi et al., 2012).

### 2. Tank experiments

#### 2.1 Exp.1: Effect of spawning activity on eDNA concentration

The orange–red variety of *O. latipes* and the wild type of *O. sakaizumii* were obtained from an aquarium shop and Fukui Prefecture (Japan), respectively. Pairs were placed in tanks in two combinations, 1) male *O. latipes* and female *O. sakaizumii* and 2) male of *O. sakaizumii* and female *O. latipes* (three tanks for each combination, dimensions 245 × 225 × 145 mm, 4 L). An aquarium without a medaka species was also prepared as the experimental control to check for cross-contamination during the aquarium experiment. The tanks were kept in the laboratory at 28°C room temperature with a 14:10 hr light/dark cycle for 3 – 10 days. To maintain optimum nutritional for the experimental fish, formula feed (Hikari medaka no esa for spawning, Kyorin corporation, Himeji, Hyogo, Japan) was provided four to five times a day.

The water was sampled on the day after spawning activity was first observed in each tank. Using a syringe, 50 mL of middle water was collected from the centre of each tank at three time points: (A) 15 min before turning on the light, (B) 15 min after turning on the light and (C) 45 min after turning on the light. The collected water samples were immediately filtered using glass fibre filters with a mesh size of 0.7 μm (GF/F, GE Healthcare Japan, Tokyo, Japan) and stored at −20°C until DNA extraction. To avoid contamination, all filtering equipment was dipped in a >10% bleach solution for >10 min, carefully washed with tap water, and finally rinsed with ultrapure water. As a filtration negative control, after all water sampling, 50 mL of middle water was collected from the experimental control and filtered, using filtering equipment in the same manner. The eDNA was extracted and quantified according to the methods described in section 4, “*Laboratory analysis*”. Spawning activity was observed between time points B and C. In all tanks, all eggs (approximately 11–23 eggs for one female) were fertilized, and thus the sperm release from a male during spawning activity was confirmed.

#### 2.2 Exp.2: Relationship between the counts of spawning activity and the increment of eDNA concentration

Ten female *O. latipes* (22.6 ± 3.6 mm standard length, mean ± SD) and six male *O. sakaizumii* (25.5 ± 1.6 mm standard length, mean ± SD) were placed in a tank (dimensions 450 × 24 × 30 mm) containing 24 L of aged tap water (aerated throughout the experiment). The tank was kept for 18 days under the same conditions as in the first tank experiment. True spawning activity was observed every day from day 5; however, spawning timing and the spawning counts were unstable.

After the pre-breeding period, water sampling was performed on days 9, 16, and 18. Spawning activity was also observed on days 10–15 and 17. However, we could not perform water sampling because of the fish’s bad condition and the observer’s schedule (S. Tsuji). All spawning activity was performed by a single pair. Using syringe, 50 mL of middle water was collected from the centre of the tank at five time points: 1, at 15 min after turning on the light, and 2–5 at approximately 1 min after the true spawning activity (from first to fourth observations). The number and timing of the spawning activity with and without egg release were recorded during water sampling (Table S1). All water sampling was performed within 1 hr and 45 min. The collected water samples were stored on ice until filtration and immediately filtered using glass fibre filters (GF/F, 47 mm; nominal pore size 0.7 μm; GE Healthcare, Little Chalfont, UK), after the water was sampled each day. On the final day (day 18), as a filtration negative control, 50 mL of ultrapure water was filtered and treated in the same manner as in the real samples. Filter samples were stored at −20°C until DNA extraction. eDNA was extracted and quantified according to the methods described in section 4. “*Laboratory analysis*”.

### 3. Field survey

The field survey was performed in each medaka species’ natural habitat to examine whether the eDNA spike, caused by spawning activity, could be observed under field conditions. The two (L01 and L02; Shiga Prefecture, Japan; *O. latipes*) and one (S01; Fukui Prefecture; *O. sakaizumii*) sampling sites were selected for each medaka species. The field type was a small water channel (width <1.0 m; L01 and S01) or a pond (<10 m in diameter; L02). We avoided publishing detailed information of each sampling site because the fish are rare species. Water sampling was performed before and after sunrise in the reproductive season (September 15 or 16, 2019) and non-reproductive season (November 9 or 10, 2019). Detailed information on sampling times, sunrise times and water temperatures are shown in Fig. 2. Water qualities were measured using a handy sensor (PCS Tester 35; Advance Instruments & Chemicals, Uttar Pradesh, India). Additionally, during the reproductive season, a visual survey was performed between sunrise and a second water sampling to confirm spawning.

**Fig 1:**
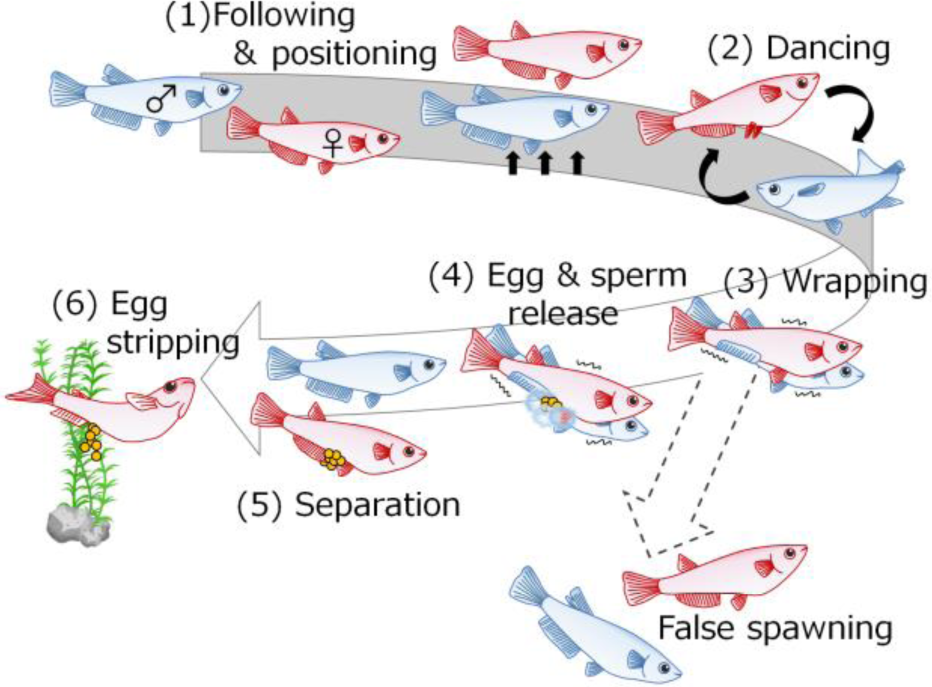
Spawning activity in medaka species.

**Fig. 2:**
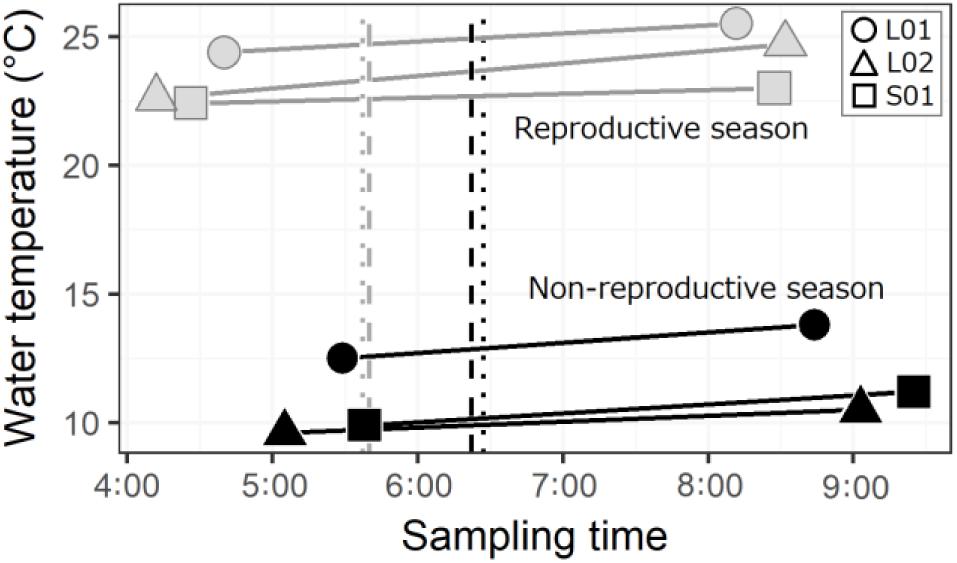
Detailed information on sampling time, sunrise time and water temperature in the field survey. Gray and black indicate a reproductive and non-reproductive season, respectively. Dashed (S01) and dotted (L01 and L02) lines indicate sunrise time.

We collected 0.5 L of surface water from each sampling site at two time points: approximately 1–2 hr before sunrise and approximately 2–3 hr after sunrise, using a disposable plastic cup (Fig. 1). The collected water sample was immediately filtered using a glass fibre filter on-site, and each filter was placed in a plastic bag and kept at −20 °C until eDNA extraction. As a filtration negative control, the same volume of ultrapure water was filtered at the end of each sampling day and treated in the same manner as the samples. eDNA was extracted and quantified according to the methods described in the section 4. “*Laboratory analysis*”.

### 4. Laboratory analysis

#### 4.1 eDNA extraction from filter samples

The eDNA extraction and purification were performed following the procedures described in Tsuji et al. (2019). Briefly, each filter sample was rolled and placed in the spin column (EZ-10; Bio Basic, Markham, Ontario, Canada) with the attached silica-gel membrane. The spin column was centrifuged for 1 min at 6,000 g to remove any excess water remaining in the filter. Then 310 µL of a mixture, composed of 200 µL of ultrapure water, 100 µL of Buffer AL, and 10 µL of proteinase K, was placed in each of the filter’s spin columns. After incubating the spin column at 56°C for 30 min, the liquid held in the filter was collected by centrifugation for 1 min at 6,000 g. To increase the eDNA yield, the remaining eDNA in the filter was recovered by adding 200 μL of Tris-EDTA buffer (pH 8.0) and centrifuging for 1 min at 6,000 g. A mixture of 100 µL of Buffer AL and 600 µL of ethanol was added to each collected liquid and mixed well by gently pipetting. The DNA mixture was subsequently purified using a DNeasy Blood and Tissue Kit (Qiagen, Hilden, Germany) following the manufacturer’s protocol, and the DNA was finally eluted in 100 µL of Buffer AE.

#### 4.2 qPCR quantification of eDNA

The number of DNA copies of each medaka species in each sample was quantified using the previously developed species-specific primer/probe sets for each medaka species (Tsuji et al. 2018) and the real-time polymerase chain reaction (qPCR) system (QuantStudio 3, Thermo Fisher Scientific, Waltham, United States). The sequence of the previously developed primer/probe sets for each medaka species is as follows: *O. latipes*: OlaND5-F (5’-TCTTTACTATAATCCTGGCAGTCCTTATC-3’), OlaND5-R (5’-CTGCTGCTAACTCTTTTTGTTGTTC-3’), OlaND5-Pr (5’-[JOE]-AATCTAACTGCTCGCAAAGTCCCACGACT -[BHQ]-3’), *O. sakaizumii*: Osa16S-F (5’-ATCTTCAAGTAGAGGTGACAGACCA-3’), Osa16S-R (5’-AACTCTCTTGATTTCTAGTCATTTGTGTC-3’), Osa16S-Pr (5’-[FAM]-TGGATAGAAGTTCAGCCTC-[NFQ]-[MGB]-3’).

The amplicon length of each primer/probe set for *O. latipes* and *O. sakaizumii* was 108 bp and 136 bp, respectively. The qPCR was conducted in 15-μL volume with a reaction solution that consisted of 900 nM of each primer (OlaND5-F/R or Osa16S-F/R), 125 nM of TaqMan probe (OlaND5-Pr or Osa16S-Pr), 0.2 μL AmpErase Uracil N-Glycosylase (Thermo Fisher Scientific), and 2.0 μL of DNA template in 1 × TaqMan Environmental Master Mix (Thermo Fisher Scientific). For each qPCR run, a four-point standard curve, with triplicate for plasmid DNA of each target region at known copies (3 × 10^1^ to 3 × 10^4^), was used to estimate the absolute eDNA concentration. Also, a PCR negative control, 2.0 μL of ultrapure water was added and analyzed instead of the DNA template to assess cross-contamination. The qPCR was performed in triplicate for all eDNA samples, standard dilution series, and the PCR negative control, with the following thermal conditions: 2 min at 50°C, 10 min at 95°C, 55 cycles of 15 s at 95°C, and 1 min at 60°C.

### 5. Statistical analysis

In tank experiment 1, the eDNA concentrations of the two medaka species were compared at the three time points using the non-parametric Kruskal-Wallis one-way ANOVA, followed by a Tukey honest significant differences (HSD) test on ranks. In tank experiment 2, the amount of spawning’s influence on the eDNA concentration was evaluated using a generalized linear mixed-effects models (GLMM; package nlme ver. 3.1– 139, Pinheiro et al. 2019) with a random effect of sampling day. In the model, the number of spawning activities and log-transformed eDNA copy numbers were set as explanatory and response variables, respectively. All statistical analyses were performed using R ver. 3. 6. 0 software (R Core Team 2019), and the minimum level of significance was set at α = 0.05.

## Results

In tank experiment 1, which examined the effect of spawning activity on eDNA concentration, the male species’ eDNA concentration increased drastically after spawning activity (time point C; 45 min after turning on the light) in all tanks regardless of species (Fig. 3A, *O. latipes p* < 0.001; Fig. 3B, *O. sakaizumii p* < 0.001). The female species’ eDNA concentration increased after spawning activity only when the *O. sakaizumii* was female (Fig. 3A, *p* < 0.05). In all tanks, the increasing rate of eDNA concentration after spawning activity was higher in males than in females. After spawning activity, male species’ eDNA concentrations were 5.9 (*O. latipes*) and 16.1 times (*O. sakaizumii*) higher, compared with those at 15 min, after turning on the light (before spawning). In both species, there were no differences in the eDNA concentrations between time point A (15 min before turning on the light) and B (15 min after turning on the light), regardless of sex.

**Fig. 3:**
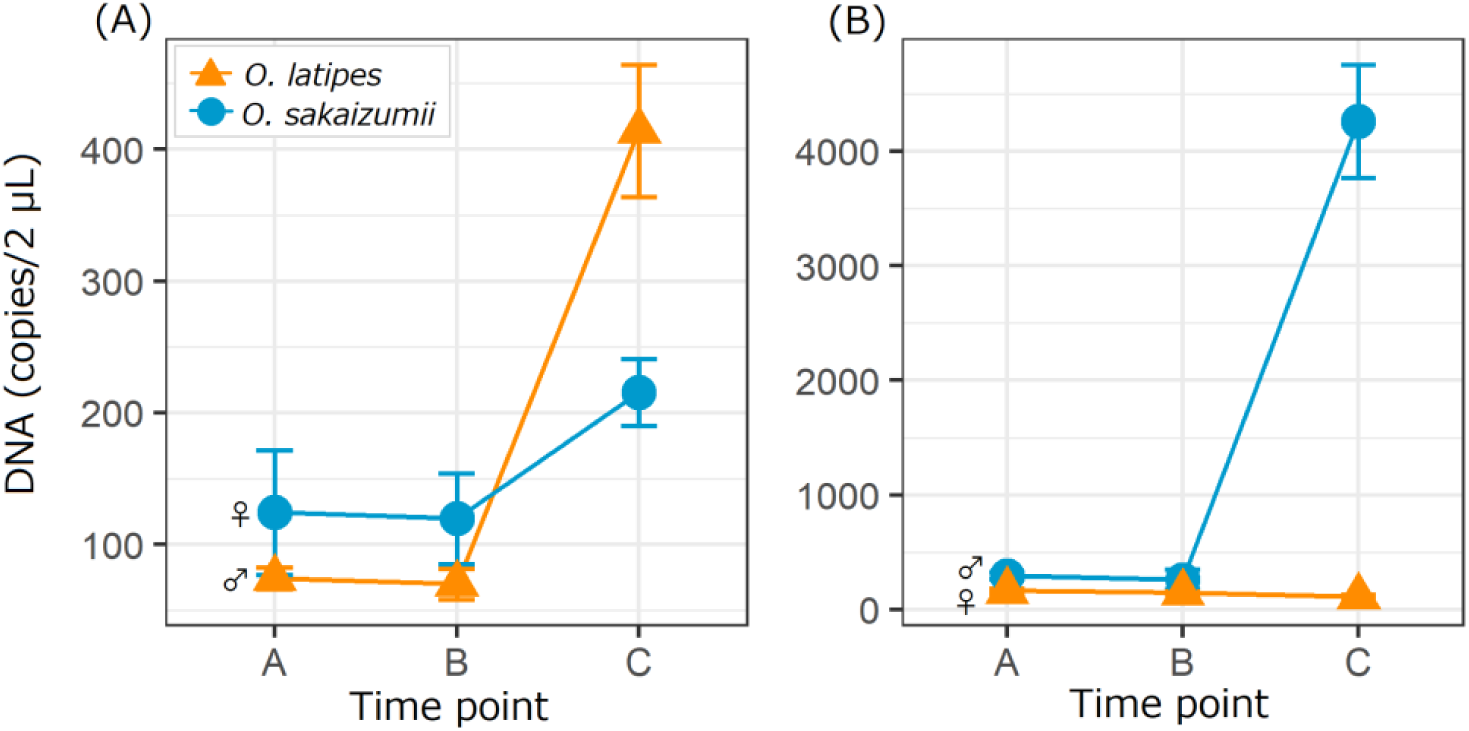
Comparison of eDNA concentration among three-time points in tank experiment 1. Three time points are as follows: A – 15 min before turning on the light; B – 15 min after turning on the light; and C – 45 min after turning on the light. Spawning activity was performed between time points B and C in all tanks. The species combination was as follows: the left panel (A) was male *O. latipes* and female *O. sakaizumii*, the right panel (B) was male *O. sakaizumii* and female *O. latipes*.

In tank experiment 2, we found a significant effect on the number of spawning activities that observed egg release (true spawning) on incremental eDNA concentration (Fig. 4A, *p* < 0.01). On the other hand, there was no significant relationship between the number of spawning activities including both true and false spawning (spawning activity without egg release) and incremental eDNA concentration (Fig. 4B, *p* = 0.06). On day 18, the first true spawning was observed within 15 minutes after turning on the light; thus, data on the period before the spawning (i.e., the number of spawning 0) could not be obtained.

**Fig. 4:**
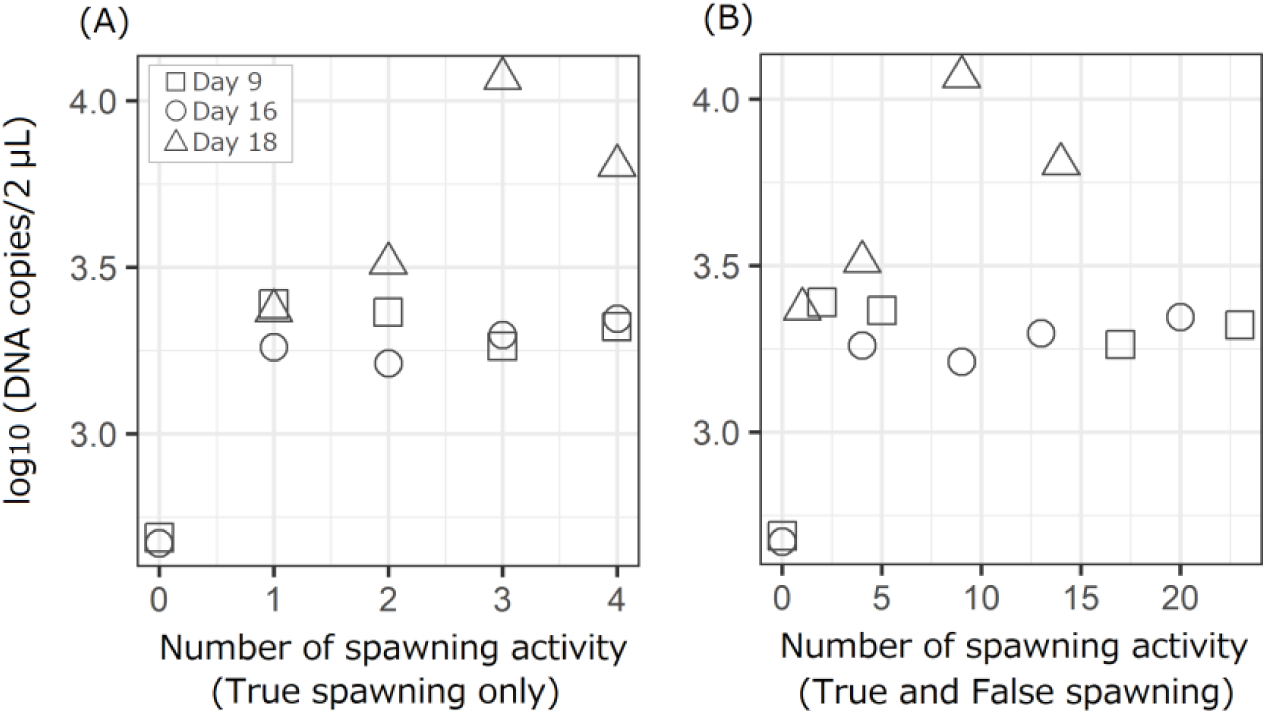
Relationship between the log10 transformed *O. sakaizumii* DNA concentrations and the amount of spawning activity (panel A, true spawning only; panel B, both true and false spawning). Squares, circles and triangles indicate sampling days 9, 16 and 18, respectively.

In the field survey, a spike in eDNA concentrations after sunrise was observed only during the reproductive season (September) in all sites (Fig. 5). The eDNA concentrations after sunrise at each site were 25.0, 7.2 and 3.1 times higher compared with this at night. Although a visual survey was performed between sunrise and the second water sampling, spawning activity was not observed. In the non-reproductive season (November), there were no notable changes in eDNA concentrations between the points before and after sunrise, regardless of species.

**Fig. 5:**
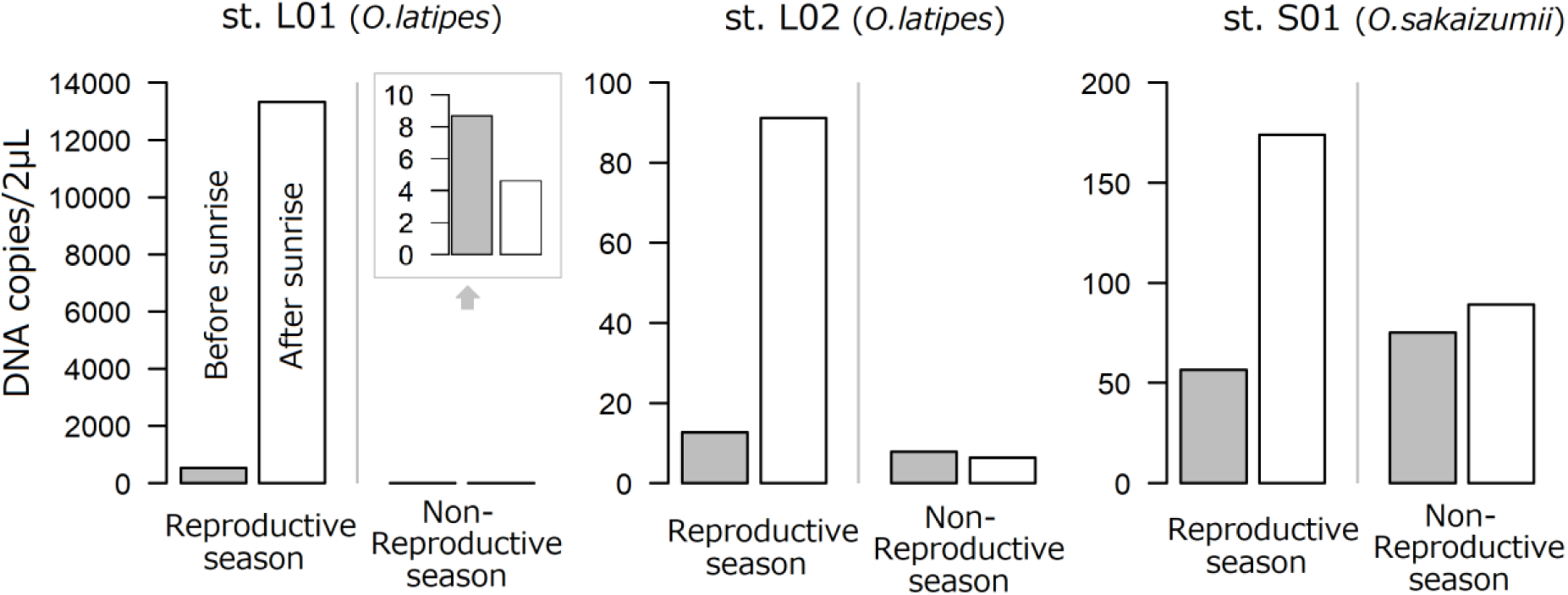
Comparison of eDNA concentration between before sunrise and after sunrise in reproductive (September) and non-reproductive (November) seasons.

In all experiments, no medaka DNA was detected in the experimental control tank, filtration negative controls and PCR negative controls, indicating that there was no cross-contamination during experiments. In all qPCR runs, the R^2^ value of the standard curve ranged from 0.984 to 0.991.

## Discussion

We found that the eDNA concentration spike was caused by spawning activities, and this can be used to monitor and understand spawning activity. Furthermore, although both sperm from males and secretions from females such as oocytes and ovarian fluid can be a source of eDNA, the significantly higher increment of eDNA on male species suggested that sperm release was the main factor of eDNA spike by spawning (Fig. 3). To our knowledge, this is the first study that experimentally identified the main factor for the increment of eDNA concentration during spawning activity. Our result is supported by the fact that the head length and head width of the sperm of *O. latipes* are 2.6 and 1.7 μm, respectively, and can be trapped by GF/F filter with a mesh size of 0.7 μm (mean) (Iwamatsu 1999). These facts also suggest that the monitoring of spawning activity based on the observation of eDNA spikes, can be applied to many other externally-fertilizing species.

Our results showed a significant positive relationship between the amount of true spawning activity with egg and sperm release and the increment of eDNA concentration, suggesting that the magnitude of eDNA spike has the potential to use for comparison of the relative amounts of spawning among sites. Since decreased data reliability due to the observation of false spawning is a serious limitation for estimating spawning amounts by conventional methods, eDNA-based monitoring which reflects only true spawning activity would be useful to assess of spawning activity. However, it is probable that the magnitude of the eDNA spike caused by spawning activity is different for each species in relation to the difference in the amount of sperm emission and/or the mating systems. Previous studies show that there is a relationship between DNA copy number and the cell abundance of artificially added sperm (Bayer et al., 2019). In tank experiment 1, we found a difference in the increase rate of eDNA caused by spawning between *O. latipes* and *O. sakaizumii*; however, the sperm cell count and observation of the amount of sperm emission were not carried out. In addition, although medaka species perform spawning activities with monogamy, there are many species which perform spawning activities by many and unspecified individuals (e.g. promis-cuity style) in other species. In this case, it remains to be seen if the relationship between the amount of spawning activity and the increment of eDNA concentration can be observed. Thus, further insights into these aspects are left future studies.

An eDNA spike after sunrise was observed in each field only during the reproductive season (September), suggesting that we can use eDNA spikes as evidence of spawning in field surveys. As a spawning occurred every day at each sampling site in the reproductive season, it seems reasonable to consider that eDNA spikes occurred every day and exhibited a diurnal change pattern. This was supported by the fact that the lowest eDNA concentration was observed before spawning on days 9 and 16 in tank experiment 2, despite a spawning activity occurring every day. Additionally, we found that the increased eDNA rates from 3.1 to 25.0 times, and eDNA was higher in the sampling site with higher water temperature. This observation could indicate that the size of the eDNA spike reflects the magnitude of spawning events in each sampling site. Especially, the water temperature was kept at optimal (i.e. approximately 25°C) in L01, the expectation was that spawning would occur most actively across the three sites. However, spawning activity was not observed by the visual survey. Observing spawning activity under field conditions is challenging because the fish are small and can easily hide in the water weeds. In our visual survey, juveniles and females with eggs were frequently observed, but not spawning activity. Our results suggest that eDNA is likely to be a sensitive tool for monitoring spawning activity and has greater success than visual surveys.

We propose that eDNA spikes may be used to monitor and understand of spawning activity. When the spawning timing is unclear, monitoring the seasonal change and/or diurnal change of eDNA concentration is an effective means of estimating spawning timing. An estimate is difficult using conventional methods such as visual survey and drift nets because these methods are strongly limited by field conditions (e.g. visibility, terrain) and the characteristics of the species (e.g. body size, infauna) and the labour-intensiveness of the tasks. We can narrow down the spawning season and time without these limitations more efficiently by observing eDNA spikes because eDNA analysis requires only water sampling at the study site. In addition, may avoid the survey restrictions on conventional spawning surveys imposed by time, labour, monitoring biases and invasiveness, for the same reason. Therefore, monitoring for eDNA concentration changes would provide promising opportunities to efficiently and non-invasively monitor spawning activity. We can also estimate the preferred spawning site by comparing the amount and/or magnitude of eDNA spikes among sites. Understanding preferred environments for spawning is critical for efficient species and/or population conservation and management. Furthermore, comparing the size of eDNA spikes can also be used to evaluate the effectiveness of artificial spawning grounds and the fish ladder as a pathway to the spawning site.

The present study also has implications for the sampling designs of future studies. When detecting rare species and/or invasive species at the initial stage of invasion, false-negative results are typically caused by low DNA concentrations (Jerde et al., 2014; Carim et al., 2019). Thus, because eDNA concentration temporarily increases, we expect to increase the detection rate with water sampling during the reproductive season and spawning hours. On the other hand, when estimating biomass by quantifying eDNA concentration, we recommend avoiding surveys during the reproductive season. The eDNA concentration will change easily and widely depending on the frequency of spawning activity, and this will lead to confusion when interpreting the results. In addition, in metabarcoding using universal primers (e.g. MiFish primers; Miya et al., 2015), the particular species DNA that increased by spawning activity is likely to inhibit PCR amplification and sequence determination of minor species’ DNA because of the consumption PCR enzyme and sequence reads. Therefore, it is necessary to design a research plan that considers whether the sampling season and hours are appropriate for the detection methods and the study purpose.

In conclusion, the eDNA spike is caused by released sperm and can be regarded as evidence of spawning. Observing an eDNA spike from spawning provides the opportunity to monitor and understand spawning timing and site with less labour, time and invasiveness than conventional methods. Knowing and understanding reproductive biology are critical and useful for efficient conservation and management of species and/or populations. Although the presented approach shows many applications as a practical tool for reproductive biology, additional studies examining the degradation and transport of released sperm would provide better insights.

## Acknowledgements

We thank K. Watanabe (Kyoto University) for providing the fish housing facility. This study was supported by Grant-in-Aid for JSPS Research Fellow Grant Number JP18J10088.

## Supporting Information

**Table S1:** The timing of all spawning activity during water sampling and egg release observation. Water sampling was carried out only when spawning activity with egg release (true spawning) had occurred.

| ID | Day 9 |  | ID | Day 16 |  | ID | Day 18 |  |
| --- | --- | --- | --- | --- | --- | --- | --- | --- |
|  | Time | Egg release |  | Time | Egg release |  | Time | Egg release |
| 1 | 0 : 46 : 50 | No | 1 | 0 : 15 : 44 | No | 1 | 0 : 3 : 12 | Yes |
| 2 | 0 : 49 : 57 | Yes | 2 | 0 : 17 : 45 | No | 2 | 0 : 7 : 19 | No |
| 3 | 0 : 59 : 7 | No | 3 | 0 : 32 : 52 | No | 3 | 0 : 7 : 35 | No |
| 4 | 0 : 59 : 58 | No | 4 | 0 : 40 : 45 | Yes | 4 | 0 : 8 : 25 | Yes |
| 5 | 1 : 4 : 38 | Yes | 5 | 0 : 49 : 23 | No | 5 | 0 : 12 : 39 | No |
| 6 | 1 : 6 : 13 | No | 6 | 0 : 52 : 39 | No | 6 | 0 : 15 : 17 | No |
| 7 | 1 : 15 : 2 | No | 7 | 0 : 56 : 18 | No | 7 | 0 : 17 : 8 | No |
| 8 | 1 : 16 : 47 | No | 8 | 0 : 57 : 15 | No | 8 | 0 : 17 : 35 | No |
| 9 | 1 : 19 : 4 | No | 9 | 0 : 57 : 51 | Yes | 9 | 0 : 18 : 49 | Yes |
| 10 | 1 : 20 : 11 | No | 10 | 1 : 4 : 36 | No | 10 | 0 : 26 : 25 | No |
| 11 | 1 : 23 : 54 | No | 11 | 1 : 6 : 36 | No | 11 | 0 : 29 : 46 | No |
| 12 | 1 : 27 : 16 | No | 12 | 1 : 7 : 23 | No | 12 | 0 : 30 : 24 | No |
| 13 | 1 : 29 : 17 | No | 13 | 1 : 8 : 27 | Yes | 13 | 0 : 31 : 17 | No |
| 14 | 1 : 31 : 56 | No | 14 | 1 : 22 : 31 | No | 14 | 0 : 33 : 9 | Yes |
| 15 | 1 : 32 : 12 | No | 15 | 1 : 22 : 54 | No |  |  |  |
| 16 | 1 : 33 : 6 | No | 16 | 1 : 23 : 37 | No |  |  |  |
| 17 | 1 : 34 : 45 | Yes | 17 | 1 : 25 : 47 | No |  |  |  |
| 18 | 1 : 39 : 48 | No | 18 | 1 : 27 : 1 | No |  |  |  |
| 19 | 1 : 39 : 58 | No | 19 | 1 : 31 : 56 | No |  |  |  |
| 20 | 1 : 41 : 54 | No | 20 | 1 : 35 : 16 | Yes |  |  |  |
| 21 | 1 : 42 : 33 | No |  |  |  |  |  |  |
| 22 | 1 : 43 : 35 | No |  |  |  |  |  |  |
| 23 | 1 : 45 : 6 | Yes |  |  |  |  |  |  |

